# Selenium-dependent metabolic reprogramming during inflammation and resolution

**DOI:** 10.1101/2021.01.16.426951

**Authors:** Arvind M. Korwar, Ayaan Hossain, Tai-Jung Lee, Ashley E. Shay, Venkatesha Basrur, Kevin Conlon, Philip B. Smith, Bradley A. Carlson, Howard M. Salis, Andrew D. Patterson, K. Sandeep Prabhu

## Abstract

Trace element selenium (Se) is incorporated as the 21^st^ amino acid, selenocysteine (Sec), into selenoproteins through tRNA^[Ser]Sec^. Selenoproteins act as gatekeepers of redox homeostasis and modulate immune function to effect anti-inflammation and resolution. However, mechanistic underpinnings involving metabolic reprogramming during inflammation and resolution remain poorly understood. Bacterial endotoxin lipopolysaccharide (LPS) activation of murine bone marrow-derived macrophages (BMDMs) cultured in the presence or absence of Se (as selenite) was used to examine temporal changes in the proteome and metabolome by multiplexed tandem mass tag-quantitative proteomics, metabolomics, and machine-learning approaches. Kinetic deltagram and clustering analysis indicated addition of Se led to extensive reprogramming of cellular metabolism upon stimulation with LPS enhancing PPP, TCA cycle, and OXPHOS, to aid in the phenotypic transition towards alternatively activated macrophages, synonymous with resolution of inflammation. Remodeling of metabolic pathways and consequent metabolic adaptation towards pro-resolving phenotypes began with Se treatment at 0 h and became most prominent around 8 h post LPS stimulation that included succinate dehydrogenase complex (Sdh), pyruvate kinase (Pkm), and sedoheptulosekinase (Shpk). Se-dependent modulation of these pathways predisposed BMDMs to preferentially increase OXPHOS to efficiently regulate inflammation and its timely resolution. Use of macrophages lacking selenoproteins, indicated that all three metabolic nodes were sensitive to selenoproteome expression. Furthermore, inhibition of Sdh with dimethylmalonate affected the pro-resolving effects of Se by increasing the resolution interval in a murine peritonitis model. In summary, our studies provide novel insights into the role of cellular Se via metabolic reprograming to facilitate anti-inflammation and proresolution.

## INTRODUCTION

Trace element selenium (Se) is incorporated as the 21^st^ amino acid, selenocysteine (Sec) via tRNA^[Ser]Sec^ (encoded by *Trsp*) dependent decoding of the UGA codon in 24 murine (25 in humans) selenoproteins (1, 2). Selenoproteins function as redox gatekeepers and catalyze reactions involving reduction of disulfides, methionine sulfoxide, and peroxides (1, 3). In addition, selenoproteins also modulate immune functions through oxidative homeostasis, prevention of iron-induced cellular ferroptosis, regeneration of reduced thioredoxin, regulation of actin repolymerization during innate immune response, and cellular calcium homeostasis (4). Although a highly debated topic, Se supplementation and its benefits in severe systemic inflammatory response syndrome, sepsis, and septic shock assumes a J-shaped curve relationship suggesting supplementation therapies primarily benefit Se-deficient patients (5, 6). However, their mechanistic role in the modulation of pathways associated with metabolic reprograming during inflammation is poorly understood (7-11).

Phenotypic plasticity of macrophages as seen in the form of classically activated M1 pro-inflammatory phenotype, upon treatment with bacterial endotoxin lipopolysaccharide (LPS), or an alternatively activated (M2) anti-inflammatory phenotype, a trait synonymous with cellular mechanisms of resolution, upon treatment with interleukin (Il)-4 or Il-13, represent two ends of the polarization spectrum (12). The resolution program involves a highly coordinated and systemic response, involving transmigration, phagocytosis, antigen presentation, and expression of pro-resolving genes such as arginase-1 and Mrc-1 (CD206) that impinge on redox-dependent signaling and cellular metabolism (12-16). Monocyte/macrophage-specific deletion of selenoproteins has led us to recognize the importance of the selenoproteome in the transition of M1 to M2-like phenotype and resolution as seen in experimental models of gut inflammation and hematologic malignancies (17-25). Recent studies from our laboratory using TMT-labeling method of non-Sec peptides in murine BMDMs indicated a temporal regulation with a general increase in the selenoproteome upon LPS stimulation in a Se-dependent manner (26). Selenow, Gpx1, Msrb1, and Selenom were highly upregulated upon stimulation with LPS when compared to other selenoproteins. Together, it appears that selenoprotein-dependent pathways of anti-inflammation and pro-resolution likely impinge on metabolic reprogramming, which is not well understood.

It is evident that changes in intracellular metabolic pathways, such as glycolysis, pentose phosphate pathway (PPP), tricarboxylic acid (TCA) cycle, fatty acid oxidation, fatty acid synthesis, and amino acid metabolism, impact cellular functions in immune cells (27, 28). Macrophages stimulated with LPS predominantly display a glycolytic metabolic phenotype (14) and decreased oxidative phosphorylation (OXPHOS), like the “Warburg effect” in cancer cells (14, 17, 27, 29). LPS-activated macrophages inhibit expression of several enzymes of TCA cycle, including succinate dehydrogenase (Sdh) complex (14) leading to accumulation of succinate and increased activation of hypoxia inducible factor-lα (Hif-1α) and IL-1β culminating in high glycolytic rates to favor M1 polarization (30). In fact, two distinct breaks have been identified in TCA cycle in M1 macrophages that include isocitrate dehydrogenase (IDH) and Sdh complex (14, 31). Interruption of Idh and Sdh results in accumulation of citrate and itaconate (32, 33), and succinate, respectively. Conversely, M2 macrophages predominantly exhibit an OXPHOS metabolic phenotype (27, 28), relying on oxidation of glucose and fatty acids to sustain OXPHOS-mediated generation of ATP. Furthermore, LPS-activation of macrophages downregulate Pkm2, while Pkm2 activation inhibit Il.-lα production as well as Hif-1α dependent genes, promoting a M2-like phenotype (34). In addition, the activity of sedoheptulosekinase (Shpk) or carbohydrate-like kinase (Carkl), which catalyzes an orphan reaction in PPP, involving the conversion of sedoheptulose to sedoheptulose 7-phosphate was shown to refocus cellular metabolism favoring an M2-like phenotype and high-redox state (35).

Using TMT labeled quantitative proteomics, targeted metabolomics, and unsupervised learning algorithms, we report here that Se in the form of selenoproteins modulate metabolic pathways involving Pkm2, Shpk, and Sdh complex to predispose activated macrophages towards OXPHOS to efficiently regulate inflammation and timely resolution. Inhibition of Sdh complex with dimethylmalonate (DMM) (36), affected the proresolving effects of Se in a zymosan model of peritonitis. Therefore, cellular Se, through its incorporation into selenoproteins, serves as a diet-derived regulator of metabolic reprograming to facilitate anti-inflammation and pro-resolution.

## RESULTS

### Activation of macrophages and effect of Se supplementation on global proteome

Based on the previously reported effects of Se and selenoproteins in favoring M2-like phenotype switching in macrophages (21), we comprehensively characterized the effects of Se on macrophage polarization following the temporal effects of LPS on differential proteome regulation in an effort to determine the underlying mechanisms by using a multi-omics approach at various time points post LPS stimulation (Figure 1). Murine bone marrow-derived macrophages (BMDMs) were polarized using LPS either in the absence or presence of Se as sodium selenite (250 nM). We first analyzed the proteomic dataset that indicated expression of prototypical LPS responsive proteins (such as Il-1β, Nos2, Cd284, Stat1, and Icam-1), in agreement with previous literature (34) (Figure S1). LPS-induced proteomic changes comprising 4710 proteins and their expression kinetics upon stimulation with LPS at 0, 4, 8 and 20 h were measured using tandem mass spectrometry and showed very high correlation (Pearson’s *r* of 0.96 and 0.99) between biological replicates (Figure 2a). Most importantly, LPS stimulation of Se supplemented BMDMs led to temporal dynamic proteome regulation (Figure 2b). Overall, ~60 % of the proteome was upregulated at 0 h, with log2 fold changes between 0 and 1.5, which eventually decreased to ~32%, 28% and 20% at 4, 8 and 20 hours post LPS stimulation, respectively. In contrast, a larger percentage of proteins remained upregulated (>0 log2 fold change) in BMDMs treated without Se, throughout all time points. However, proteome upregulation (>1.5 log2 fold change) was observed only in cells treated with Se (Figure 2b).

**Figure 1.**
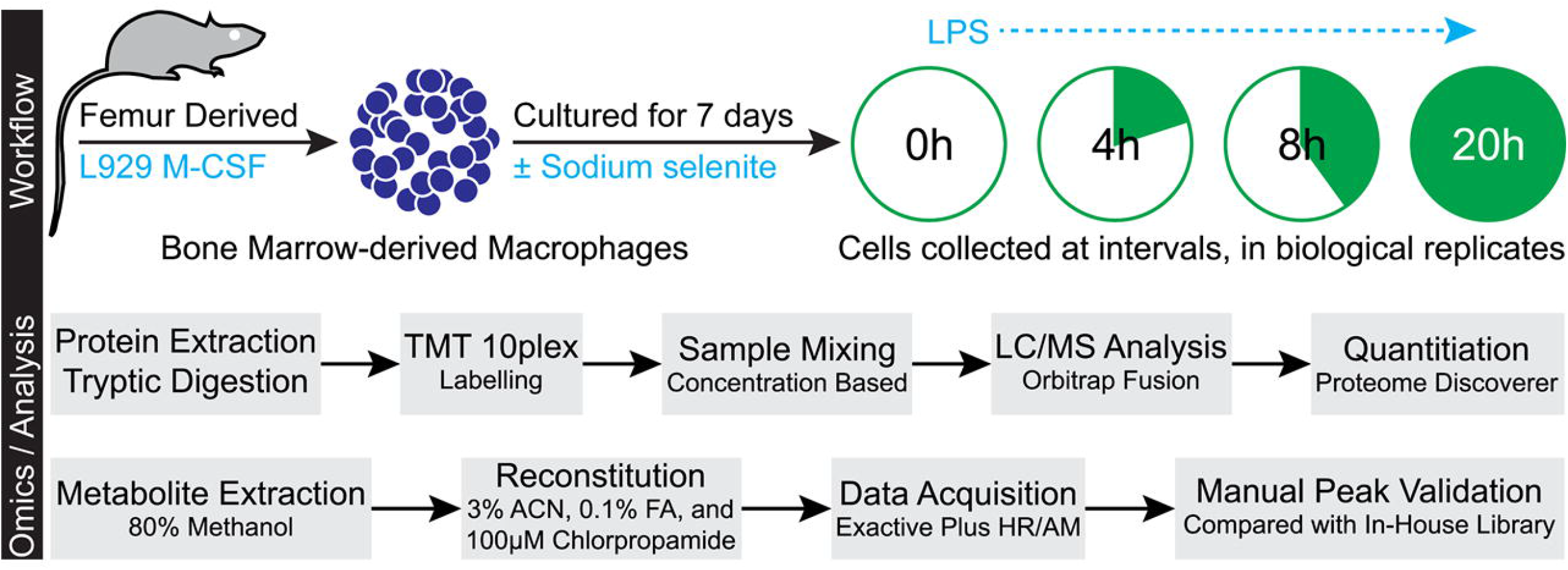
Experimental design and data acquisition. Bone marrow-derived macrophages (BMDMs) were harvested and cultured for seven days either in presence or absence of Se (as sodium selenite, Na_2_SeO_3_, 250 nM). BMDMs were stimulated with 100 ng of LPS for indicated time periods up to a maximum of 20 h. Cells were harvested following two washes with PBS and lysed to extract the proteome. Proteome was extracted and subjected to tryptic digestion followed by TMT labeling. Datasets were acquired on Lumos Fusion MS and analyzed by using Proteome Discoverer. Metabolites were extracted in methanol, dried, and reconstituted in mobile phase with chlorpropamide as internal standard. Dataset was acquired on Exactive Plus MS. Corresponding metabolite ion chromatograms were extracted using Xcalibur and manually inspected.

**Figure 2.**
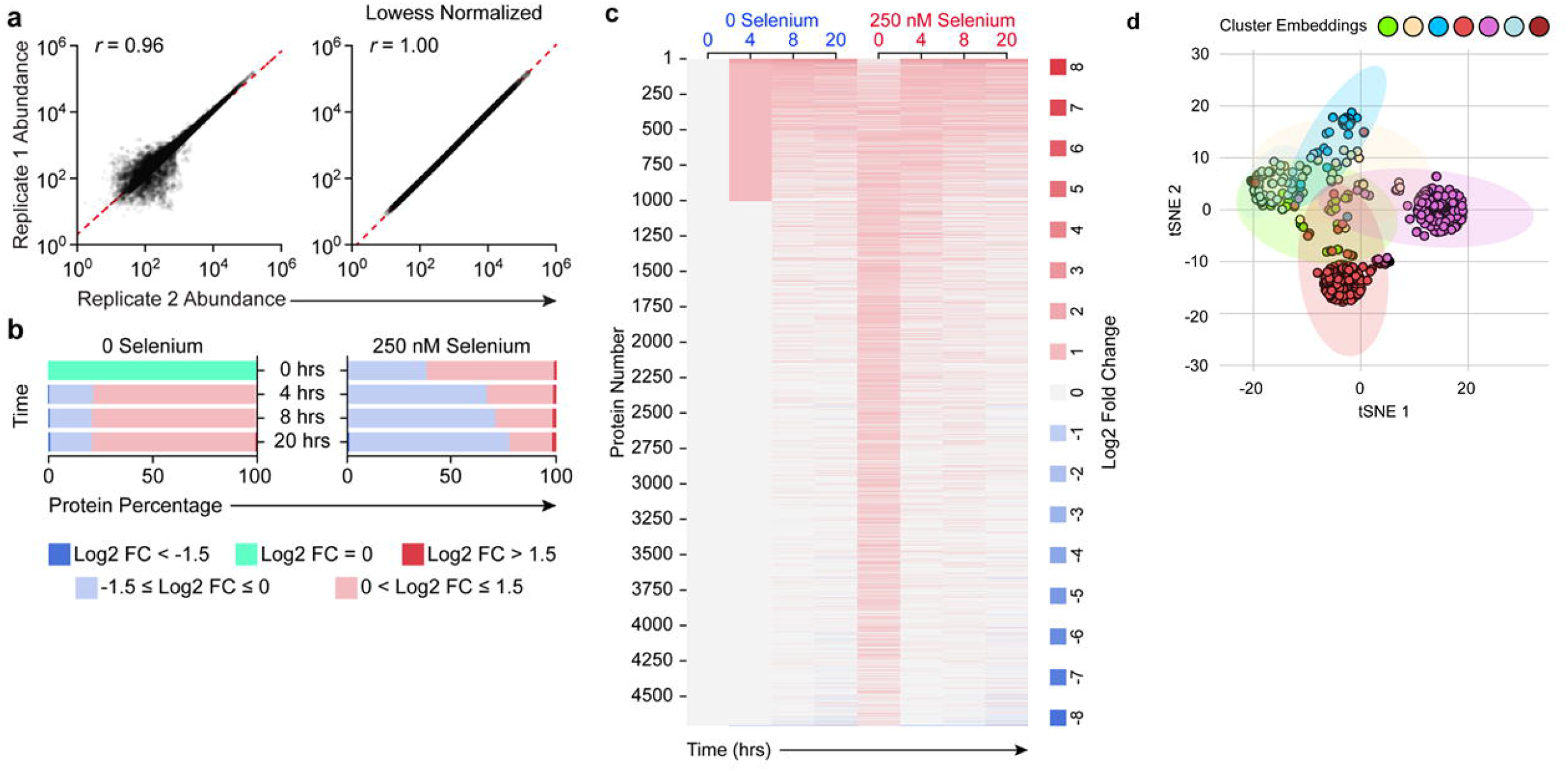
Supplementation of macrophages with Se effects global proteomic changes upon treatment with LPS. BMDMs were stimulated with LPS for indicated time periods up to a maximum of 20 h. Cells were harvested and lysed to extract the proteome. Proteomic analysis is described in methods. BMDMs cultured in Se-deficient media and not subjected to any stimulation were used for comparison. Data was normalized to β-actin and abundance values are expressed as relative to zero-hour zero-selenium cells (naïve cells). **a.** Scatterplots show biological replicates for post LPS are well correlated (Pearson’s r of 0.96). **b.** Bar chart show the overall proteome changes during post LPS stimulation. Values are represented as log2 fold changes. **c.** Heatmap showing the modulation of global BMDM proteome post LPS stimulation. **d.** The t-SNE analysis and k-means clustering of proteomic changes post LPS show seven distinct clusters, respectively.

By applying clustering analysis of proteins with similar kinetics and magnitude of induction (Figure 2c), we identified 121 proteins that were differentially regulated at 0 h in Se treated cells compared to unstimulated Se-deficient BMDMs (“naïve” cells). While majority of the selenoproteins were increased as reported earlier upon exogenous Se addition (26), nuclear factor-erythroid 2-related factor 2 (Nrf2) target genes, such as Txnrd1 (thioredoxin reductase1) and Txnrd2, and Hmox1 (hemeoxygenase 1) were also increased by Se-supplementation followed by LPS stimulation for 8 h. In contrast, 510 proteins were differentially regulated upon stimulation with LPS for 20 h (Figure 2c). Interestingly, clustering analysis indicated heterogeneity within LPS responsive proteins altered upon Se supplementation that separated into seven distinct clusters (Figure 2d; Figure S2). Specifically, clusters 1, 3, 4, 5 and 6 were enriched with 289, 631, 1292, 1496 and 536 proteins, respectively, upon LPS treatment (Figure S2). Cluster 1 represented early induced proteins impacted between 4 to 8 h post LPS treatment. The magnitude of protein induction was higher in the presence of Se compared to 0 Se, which included proteins involved in TCA cycle and electron-transport chain (ETC) that were up-regulated that was further confirmed through pathway analysis. Significant enrichment of TCA cycle and OXPHOS pathways in clusters 1, 5 and 6 showed induction of pathway-associated proteins from 4 h post LPS stimulation. Interestingly, we observed maximally expressed proteins in cluster 1, which corroborated the upregulation of proteins associated with TCA cycle and OXPHOS (Figure S2). Our analysis indicated that remodeling of metabolic pathways and consequent metabolic adaptation towards pro-resolving phenotypes started with Se treatment of Se-deficient BMDMs in the absence of any stimulation and became most prominent around 8 h post LPS stimulation.

### Se-dependent modulation of enzymes and metabolites in TCA cycle, glycolysis, and PPP

Initiation and resolution of inflammation are energy demanding processes that require timely adaptation of cellular metabolism for support. Since proteomic analysis suggested a plausible metabolic alteration in response to Se treatment, we examined the metabolic consequences of Se supplementation on macrophage activation.

Intracellular metabolites were examined in BMDMs cultured either in the presence or absence of Se followed by stimulation with LPS for 2, 4, 8 and 20 h. LPS treatment profoundly affected a wide variety of intracellular metabolites as depicted in Figure 3a. t-SNE analysis indicated that LPS treatment groups were notably separated from each other (Figure 3b). An interesting trend was observed for all time points post LPS treatment in cells cultured in the absence and presence of Se, where 0 to 8 h intervals were closest to each other. However, they were also most separated with respect to presence or absence of Se. This difference was more profound at 20 h post LPS treatment between the two groups, further emphasizing the contribution of Se in reprograming of metabolites involved in TCA cycle, glycolysis, PPP, energy molecules, redox molecules and amino acids implicated in resolution of inflammation (Figure 3a). Based on these studies, a targeted metabolomics approach was adopted to further investigate and compare the association between TCA cycle, glycolysis, and PPP, upon stimulation with LPS.

**Figure 3.**
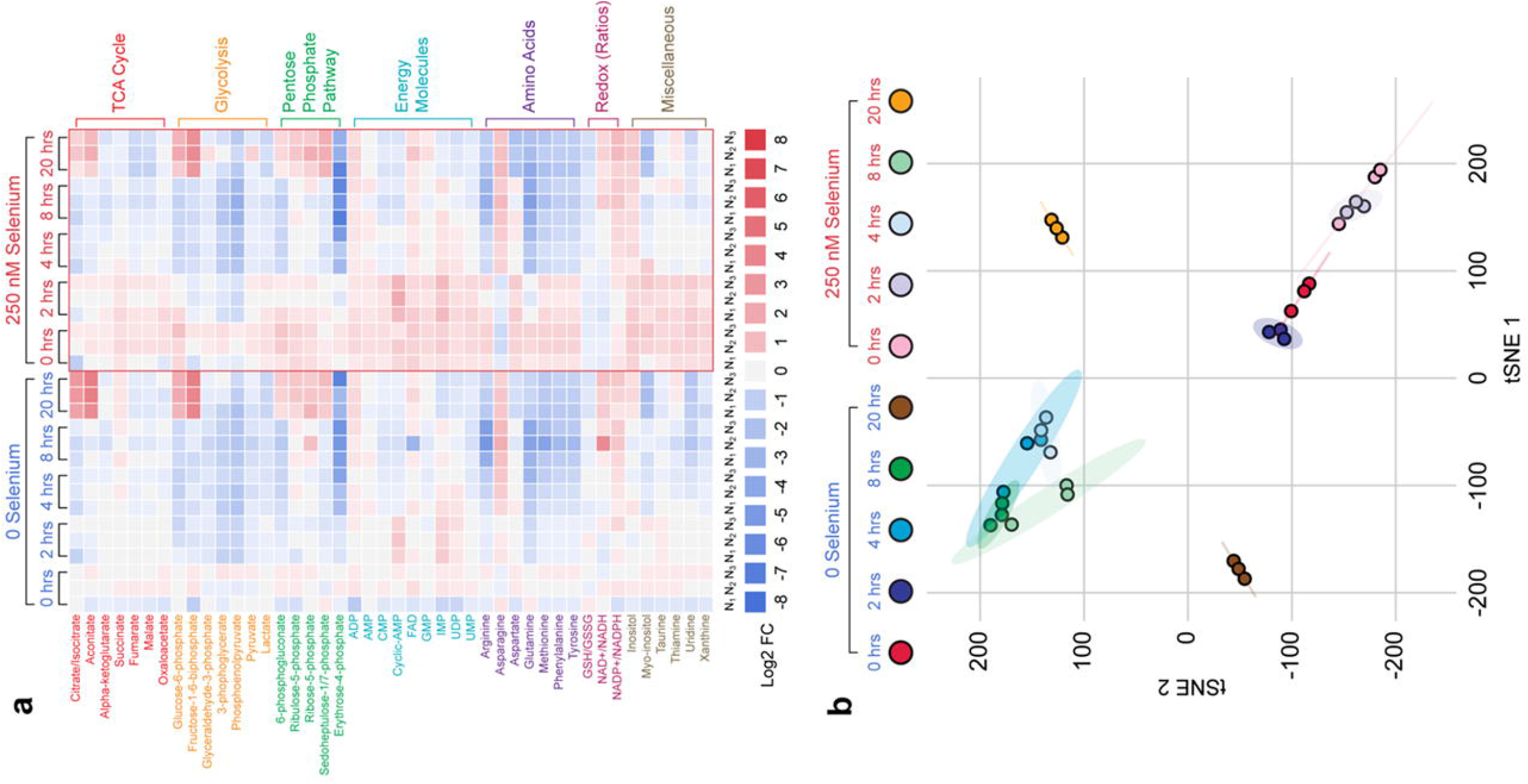
Differential modulation of key metabolites via Se in LPS-stimulated BMDMs indicate a Se-dependent reduction in glycolytic pathway and increased TCA cycle. Metabolites were extracted in methanol, dried, and reconstituted in mobile phase aqueous methanol containing 100 μM chlorpropamide as internal standard. All samples were acquired in biological triplicate and randomized before 10 μl was acquired in LC-MS using reverse-phase UHPLC coupled to an Exactive Plus orbitrap MS. A total of 47 metabolites were profiled in biological triplicates. Individual metabolite extracted-ion chromatograms were used, with peak area determined. All peak areas were normalized to internal standard chlorpropamide and abundance was calculated relative to that of zero-hour zero-selenium cells (naïve cells). **a.** Heat map profiling of inflamed BMDMs show log2 fold change values of all 47 metabolites. All displayed log2 fold changes are represented in biological triplicates. **b.** The t-SNE method applied to all samples show distinct clusters separating the samples by time post LPS stimulation.

We first examined citrate/isocitrate levels in BMDMs cultured with and without Se, before and after LPS treatment. Untreated cells failed to show any temporal differences as in cells stimulated with LPS for the first 8 h. However, prolonged treatment for 20 h with LPS significantly increased citrate/isocitrate levels in Se-deficient cells when compared to macrophages treated with Se (Figures 3a and 4). Additionally, aconitate modulation followed a similar trend, with an increase at 20 h post LPS in Se-deficient cells (Figures 3a and 4) suggesting a constant flux of citrate through the TCA cycle. Temporal modulation of succinate in response to exogenous Se treatment before and after LPS stimulation was interesting. Succinate levels were high in unstimulated cells in the presence of Se (compared to those without Se) that showed a gradual decline upon LPS stimulation starting at 4 h that persisted even at 20 h (Figures 3a and 4). On the other hand, succinate levels increased in Se-deficient cells gradually over the 20 h post LPS treatment. Furthermore, fumarate, a product of succinate oxidation, and its hydrated product, malate, followed an identical pattern to that of succinate (Figure 4). Interestingly, LPS-dependent temporal decrease in succinate, fumarate, and malate was not observed in BMDMs derived from *Trsp^fl/fl^LysM^Cre^ mice*, despite the presence of 250 nM selenite (Figure 4) clearly implicating a unique role for Se/selenoproteins in ensuring an uninterrupted TCA cycle.

**Figure 4.**
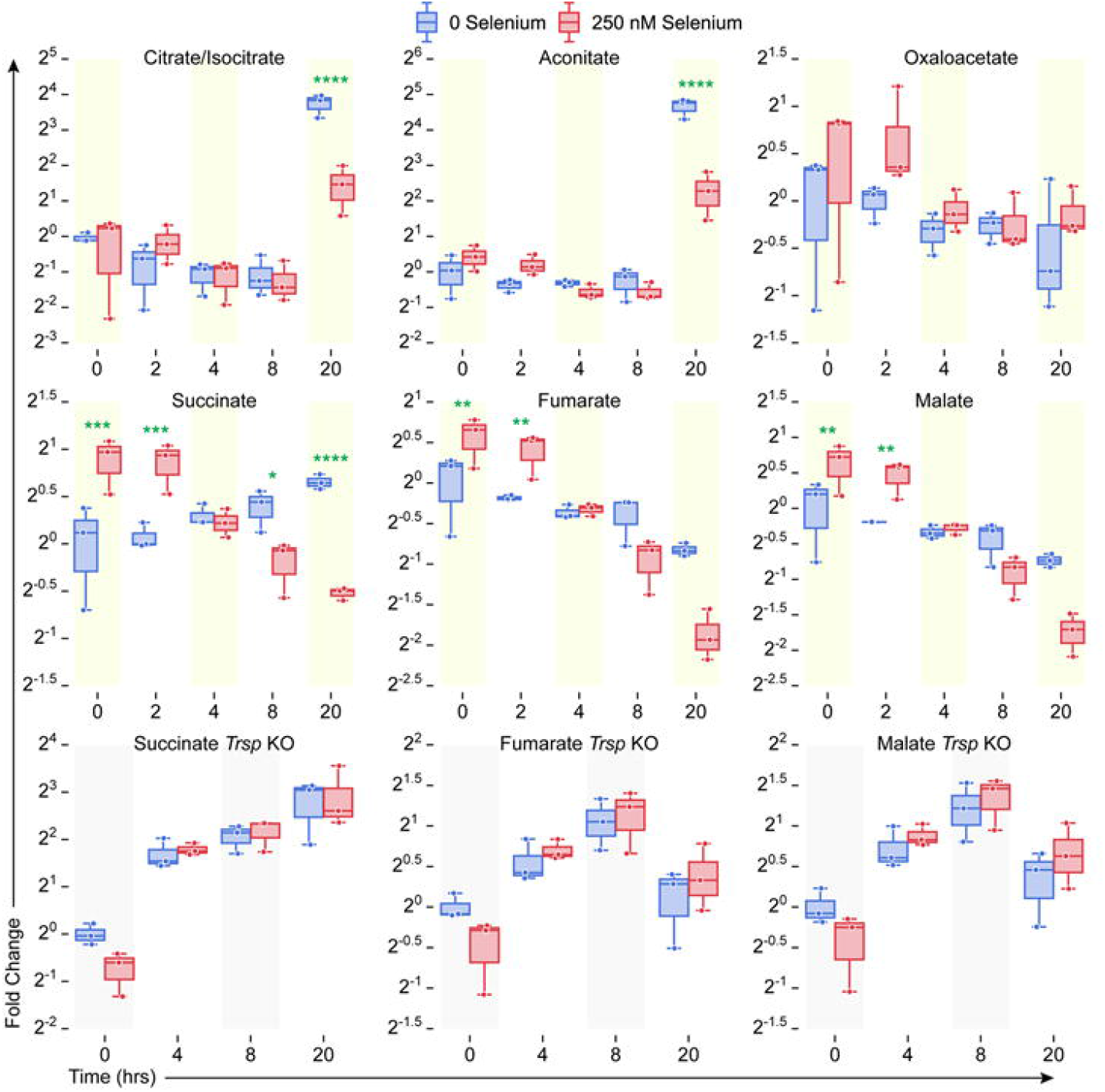
Selenoprotein-dependent changes in TCA cycle metabolites in WT and *Trsp^fl/fl^ LysM^Cre^* BMDMs post LPS stimulation. Metabolites were extracted from BMDM cells isolated from WT and *Trsp*^fl/fl^LysM^Cre^ mice as described earlier and analyzed by LC-MS. BMDMs from (n= 3) mice per genotype were cultured as described earlier and stimulated for various time points 0-20 h with LPS. Metabolite peak areas were normalized to internal standard chlorpropamide and abundance was calculated relative to that of zero-hour zero-selenium cells (naïve cells). Se supplementation of WT BMDMs with 250 nM Se (as selenite) showed dramatic regulation of succinate levels post LPS stimulation when compared to the BMDMs isolated from *Trsp^fl/fl^LysM^Cre^* mice.

Given the Se-dependent decrease in succinate, we further examined the modulation of proteins associated with TCA cycle. Notably, protein levels of isocitrate dehydrogenase and individual subunits of the Sdh complex were higher in Se-supplemented BMDMs (Figure 5). The mitochondrial proteome was classified into six clusters based on the kinetic regulation of protein expression upon LPS stimulation (Figure S3). Western blot analysis of Sdha (flavoprotein subunit of complex II) corroborated proteomic dataset showing higher levels of expression at 0, 4, and 8 h post LPS stimulation (Figures 5a-c). However, expression of Sdha in *Trsp* KO BMDMs was decreased at time points 0, 4, and 8 h post LPS, but increased at 20 h in the presence of 250 nM selenite treatment indicating an opposite effect that corroborated the modulation of cellular succinate levels (Figure 5b and c). Western blot analysis of the expression of Sdhb (iron-sulfur subunit of complex) was higher in the WT BMDMs compared to BMDMs of *Trsp* KOs cultured with selenite (Figures 5b and d). In addition, fumarate hydratase (fumarase; Fh1) also increased in LPS-stimulated BMDMs cultured with Se, while malate dehydrogenase 2 (Mdh2) displayed an increase at 4 h post LPS stimulation in Se-supplemented BMDMs compared to the Se-deficient counterpart (Figure 5a). Interestingly, flow cytometry-based ROS measurements indicated cytosolic ROS levels were higher in Se-deficient cells that decreased after a transient increase with LPS stimulation followed by an increase at 20 h compared to the Se-supplemented cells where the levels were held steady (Figure S4a). However, Se-supplemented cells displayed higher levels of mitochondrial ROS (predominantly superoxide) (Figure S4b), suggestive of an active TCA cycle and OXPHOS, which corroborates our proteomic and metabolomic studies.

**Figure 5.**
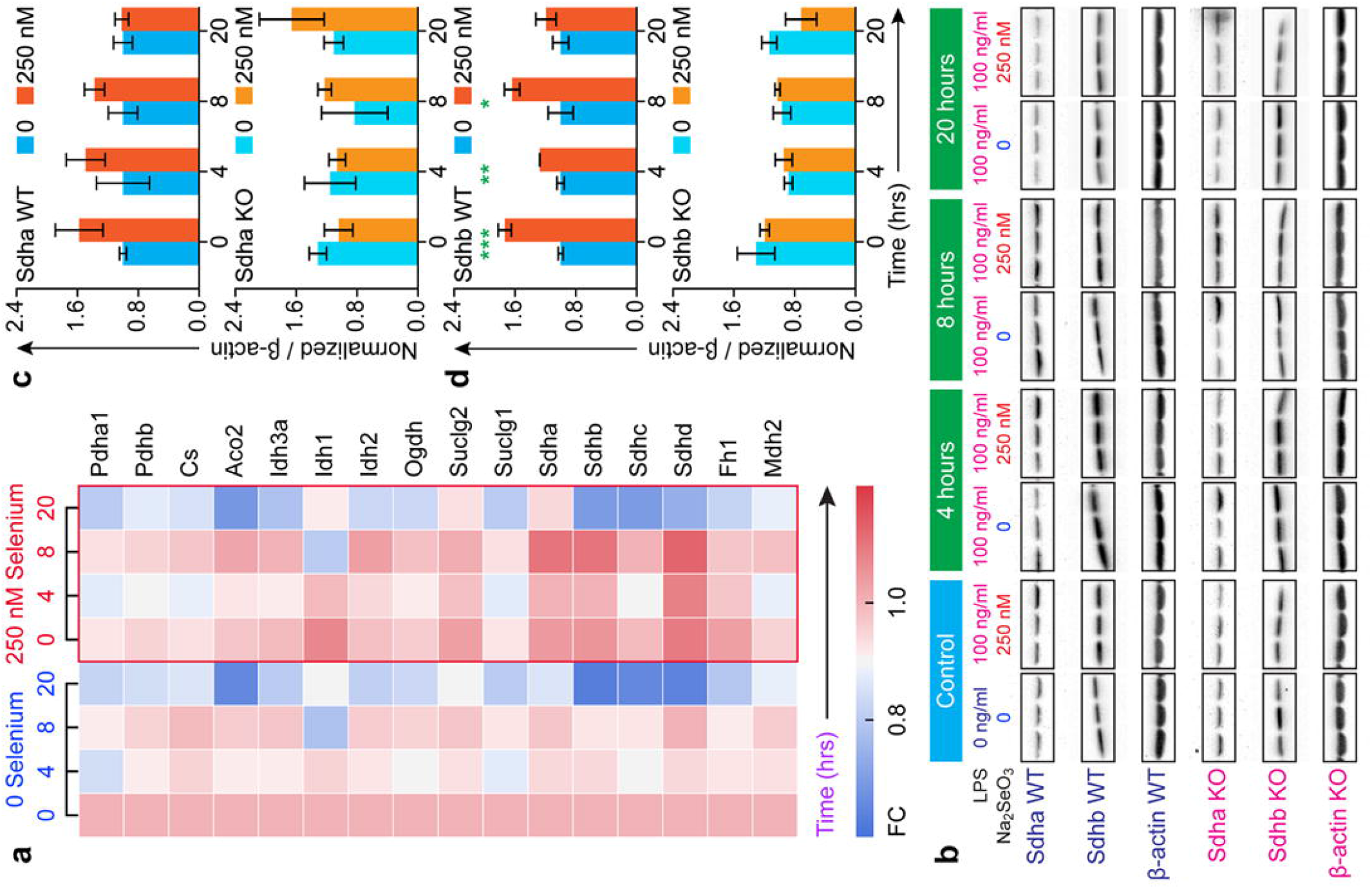
Selenoprotein-dependent changes in TCA cycle proteins. **a.** Heat map showing temporal regulation of TCA cycle proteins as determined by proteomic studies. **b.** Western blot of Sdha and Sdhb. BMDMs were isolated from wild type (WT) and *Trsp^fl/fl^ LysM^Cre^* mice respectively and incubated with L929 containing DMEM without Na_2_SeO_3_ (250 nM Se) for eight days. Cells were stimulated with 100 ng LPS on day eight for one hour and cells were harvested at 4, 8, and 20 hours, followed by protein extraction and western blot analysis of BMDMs in biological triplicate per genotype. Densitometric analysis of Sdha and Sdhb expression was performed using Image J and normalized to that of β-actin. **c** and **d**. Densitometric evaluation of western immunoblots for the expression of Sdha and Sdhb in BMDM incubated with Se compared to those cultured without Se before and after LPS stimulation. Representative data shown is mean ± SEM of n =3 per genotype at each time point post LPS treatment. *p<0.05; **p<0.005; ***p<0.0005.

To further examine if Se supplementation of BMDMs impacted key glycolytic pathway metabolites before or after LPS stimulation, we examined the temporal modulation of the following metabolites: glucose 6-phosphate (G6P), fructose 1,6-bisphosphate (F16BP), glyceraldehyde 3-phosphate, 3-phosphoglycerate, phosphoenolpyruvate, pyruvate, and lactate. These glycolytic metabolites showed varying levels of modulation in Se-deficient and Se-supplemented cells before and after LPS stimulation. Levels of G6P, glyceraldehyde-3-phosphate, 3-phosphoglycerate, pyruvate, and lactate were higher in Se-supplemented macrophages compared to their Se-deficient counterparts prior to stimulation. Upon stimulation with LPS, a time-dependent decrease in these metabolites were observed in Se-supplemented cells, especially at 8 h post LPS (Figure 3a, Figure S5). In contrast to WT BMDMs, unstimulated BMDMs isolated from *Trsp* KO mice showed low levels of phosphoenolpyruvate and pyruvate that remained unchanged upon addition of sodium selenite (250 nM). LPS-stimulation of these BMDMs lacking selenoproteins showed a markedly different temporal regulation of these metabolites, in contrast to WT BMDMs, highlighting an important role for selenoproteins in cellular metabolism during inflammation (Figure S5). Furthermore, proteomic expression profile of glycolytic enzymes showed generally reduced levels in Se-supplemented BMDMs compared to the Se-deficient BMDMs before and/or following stimulation with LPS (Figure 6a). Gpi, Pfk-l, Aldoa, Aldoc, Tim, Gapdh, Pgk1, Pgm2, Eno1, and Pkm were all decreased at 8 h post LPS stimulation in Se-supplemented BMDMs compared to their Se-deficient counterparts (Figure 6a). Particularly, Eno1 and Pkm (Pkm2) were drastically decreased in Se-supplemented BMDMs post 8 h LPS stimulation. The phosphorylated (and dimeric) form of PKM exhibited an increase at 0, 4 and 8 h post LPS treatment in the presence of Se, while an opposite effect was seen in the absence of Se. Interestingly, *Trsp* KO BMDMs cultured in the presence of Se (250 nM) showed similar trends to the Se-deficient WT BMDMs with regard to phosphorylated Pkm2 (Figure 6b and c).

**Figure 6.**
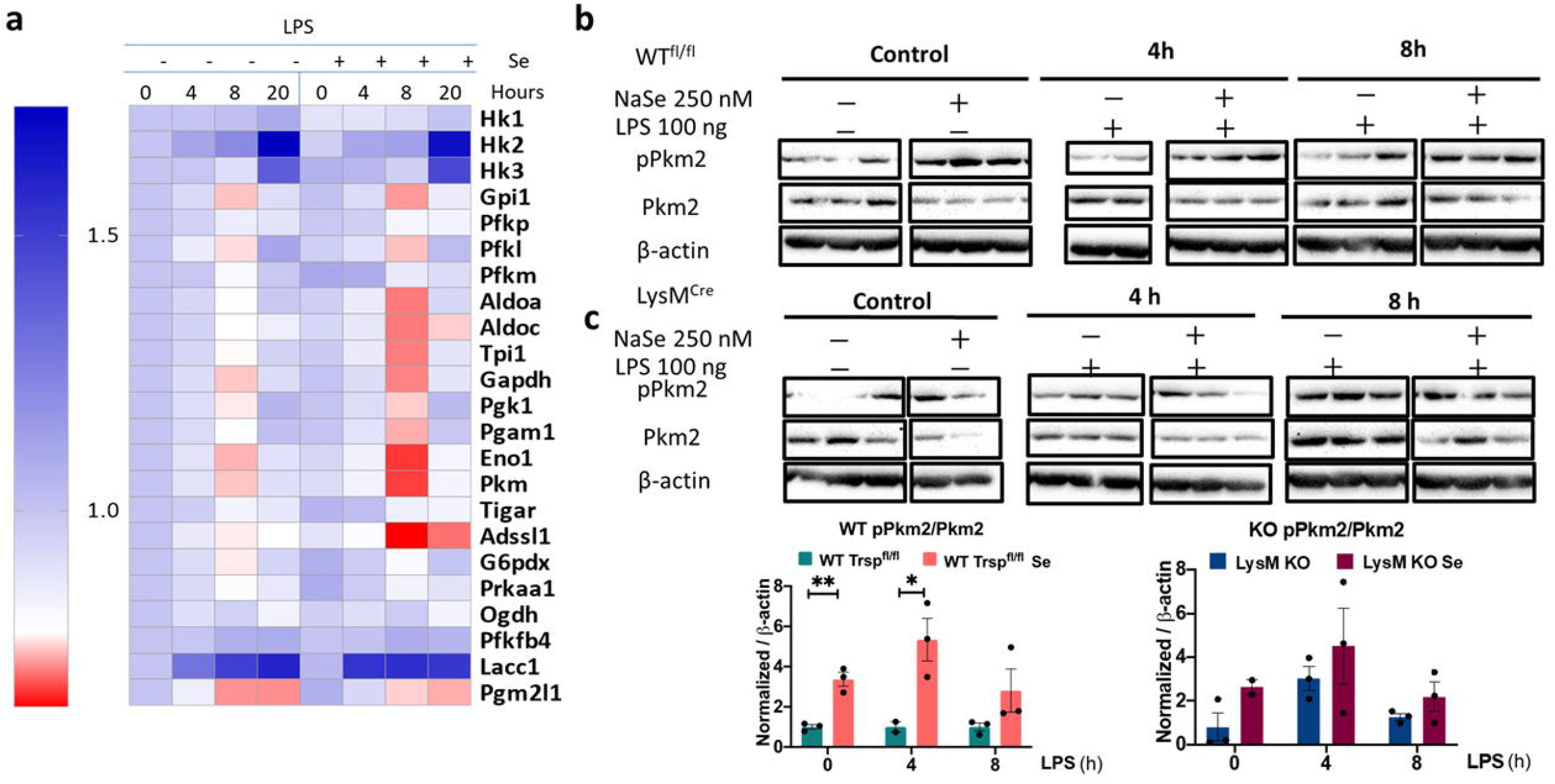
Selenoprotein-dependent changes in glycolytic cycle proteins. **a.** Heat map showing temporal changes in glycolytic pathway proteins as determined by proteomic studies. **b and c**, Western blot of pPkm2 and Pkm2 (total). BMDMs were isolated from wild type (WT) and *Trsp^fl/fl^ LysM^Cre^* mice. Cells were stimulated with 100 ng LPS on day eight for one hour and harvested at 4, 8, 20 hours, followed by protein extraction and western blot analysis. Densitometric analysis of western immunoblots using Image J program provided pPkm2:Pkm2 ratio that was normalized to β-actin and compared to cells cultured without Se. Representative data shown is mean ± SEM of n =3 per genotype at each time point post LPS stimulation. *p<0.05; **p<0.005.

Key PPP pathway metabolites were analyzed to examine the impact of changes in cellular Se status before or after LPS stimulation. 6-phosphogluconate, ribulose 5-phosphate, ribose 5-phosphate, and erythrose 4-phosphate levels decreased upon Se supplementation, especially at 8 h post-LPS treatment (Figures 3a, Figure S6). Interestingly, Shpk (Carkl), a key enzyme that catalyzes the formation of sedaheptulose-7-phosphate, was downregulated in Se-deficient BMDMs when compared to those cultured in the presence of Se. Accordingly, sedoheptulose 7-phosphate was higher in Se supplemented BMDMs, particularly at 0-2 h, and later at 20 h post LPS treatment, suggesting that Se affects expression regulation of Shpk (Carkl) (Figures 3a, Figure S6). Western blot analysis of Shpk expression indicated modulation in WT BMDMs treated with Se starting at 4 h with differences persistent up to 20 h post LPS treatment. Such modulations were absent in *Trsp* KO BMDMs, suggesting that the effect of Se is mediated through selenoproteins (Figures 7a and b). Collectively, our results lend credence to the hypothesis that Se supplementation of BMDMs increases mobilization of metabolites via PPP to potentially assist in various cellular functions to effect resolution, a thermodynamically active process.

**Figure 7.**
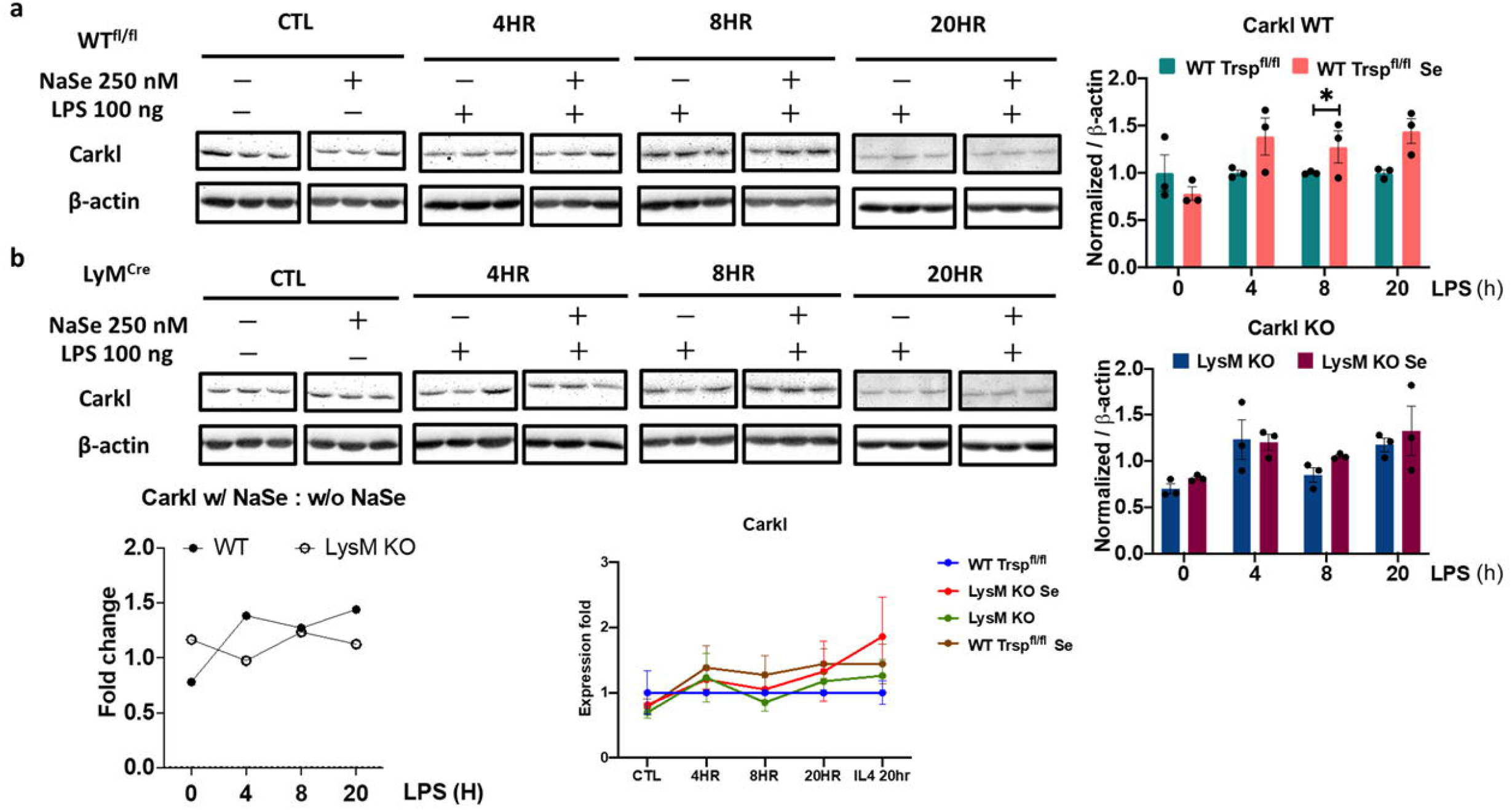
Selenoprotein-dependent changes in Shpk (Carkl). **a and b**, Western immunoblot of Carkl expression upon stimulation of BMDMs with LPS in the presence or absence of Se. BMDMs were isolated from wide type (WT) and *Trsp^fl/fl^ LysM*^Cre^ mice and cells were stimulated with 100 ng LPS on day eight for one hour and harvested at 4, 8, 20 hours, followed by protein extraction and western blot analysis. Densitometric analysis using Image J of Shpk (Carkl) expression that was normalized to the expression of β-actin. Shpk fold change in BMDMs incubated with Se were compared to those without Se. Representative data shown is mean ± SEM of n =3 per genotype at each time point post LPS stimulation. *p<0.05; **p<0.005.

### DMM treatment negatively impacts Se dependent resolution of inflammation

Given that our data indicated Se supplementation led to decreased succinate levels in addition to the modulation of Sdh expression, we examined if inhibition of succinate oxidation in mice negatively impacted the effect of Se in the resolution of zymosan-induced peritonitis. Mice on Se-supplemented diet (0.4 ppm) were treated intraperitoneally with either vehicle control (PBS) or DMM (160 mg/kg) in PBS, a competitive inhibitor of Sdh, 3 h prior to Zymosan A injection in a resolution indices (R_i_) experiment (see Methods). DMM treatment followed by Zymosan A activation led to a higher percentage of neutrophils (~75%) with a R_i_ = ~36 hours, whereas PBS-treated mice had ~40% neutrophils and an R_i_ = ~15 hours (Figure 8a). DMM treatment also led to a slight increase in total F4/80^+^ macrophages in response to zymosan A stimulation (Figure 8b), while decreasing the M2-like macrophages (F4/80^+^ CD206^+^) in the peritoneal lavage fluid (Figure 8c) compared to the PBS control. These studies further corroborate *in-vitro* observations and suggest that a Se-dependent increase in succinate oxidation is key to timely resolution of inflammation.

**Figure 8.**
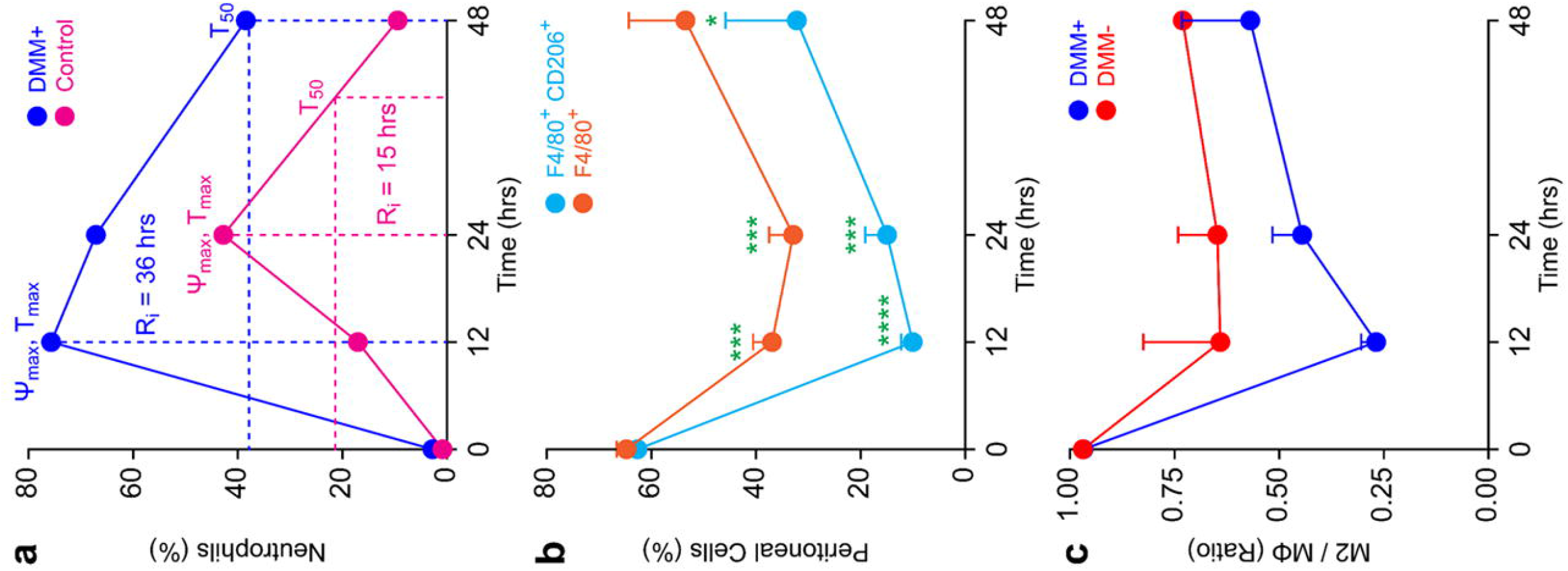
DMM treatment of Se-Supplemented mice decreases the resolution index in a zymosan-induced peritonitis model. Se-Supplemented mice were treated intraperitoneally with or without DMM in PBS (160 mg/kg of body weight) or PBS three hours prior to administration of Zymosan A (10 mg/kg body weight) intraperitoneally. Mice were euthanized at 0, 12, 24, and 48 hours post-Zymosan A injection, and peritoneal exudates were collected, centrifuged at 400× g for 5 minutes at 4 °C, washed with flow buffer (PBS containing 2% FBS and 100 I.U. penicillin and 100 μg/mL streptomycin), resuspended in 1 mL flow buffer, counted by trypan blue, and 100,000 viable cells were analyzed by flow cytometry. **a.** Resolution index (R_i_) calculation based on percentage of neutrophils (Ly6-G+ cells) in the peritoneal lavage fluid at each time point. Ψ_max_ and T_max_ represent the highest percentage of cells post zymosan treatment and the time point at which it occurs, respectively. Data shown are representative of biological triplicates. **b.** Percentage of total macrophages (F4/80^+^) and M2-like macrophages (F4/80^+^ CD206^+^) in peritoneal lavage fluid as a function of time post Zymosan A treatment. **c.** Ratio of the total macrophages to M2-like macrophages in the peritoneal lavage fluid as a function of time post Zymosan A treatment. Data shown are mean ± SEM of n = 3-4 per treatment group; ***p<0.0005. ****p<0.0001.

## DISCUSSION

Resolution of inflammation is an active and highly orchestrated process that restores normal functional tissue homeostasis (37), where macrophages play an integral role (11, 12). Plasticity of these cells arises from their ability to skew metabolism from glycolysis and fatty acid synthesis in M1 macrophages to OXPHOS and fatty acid β-oxidation, in M2 macrophages, to meet their energy demands for survival and functionality (27). Here, using multi-omics platforms we examined the effect of micronutrient Se and selenoproteins on the reprogramming of metabolic pathways involving Pkm, Shpk (Carkl), and Sdh complex to predispose activated macrophages towards OXPHOS and eventually facilitate resolution of LPS-induced inflammation.

In addition to increased mitochondrial ROS levels, significant reduction in both citrate and aconitate, along with increased Idh expression, decreased α-ketoglutarate, succinate, fumarate, malate, and oxaloacetate by exogenous Se suggested increased mobilization of TCA cycle, when compared to Se-deficient cells. Temporal increase in succinate post LPS, which moonlights as a key mediator of inflammation (30), was reversed by Se with an increase in fumarate and malate. Inhibition of Sdh complex with dimethylmalonate (DMM) (36), affected the pro-resolving effects of Se in a zymosan-induced peritonitis in mice. Though the exact mechanisms of regulation of Sdh complex by selenoproteins are unclear, inhibition of succination of reactive cysteine in fumarate hydratase (38) by Se could serve as a potential mechanism to help restore the broken TCA cycle.

Surprisingly, despite the increase in succinate that is known to increase Hif-1α, the expression of Hif-1α or its characteristic downstream genes, including Il-1β, glycolytic enzymes, and glucose transporters (30), was not observed in Se-deficient cells. However, we observed a decrease in glycolytic enzymes in macrophages supplemented with Se, suggesting considerable reduction in glycolytic rates. Pkm, a key regulatory node, was found to be significantly down regulated in macrophages supplemented with Se. Pkm is also regulated by phosphorylation status by upstream kinases that is indicative of dimer/tetramer formation (39). The dimeric and phosphorylated form of Pkm2, which is enzymatically inactive, also positively regulates its expression and that of Hif-lα-dependent genes (40). The effects we observed was not similar to that reported with DASA-58, a selective activator of Pkm2, which inhibited the Hif-1α-dependent induction of glycolytic proteins by LPS (34), suggesting Hif-1α-independent control of metabolic effects in the presence of Se. Along with PKM, there was a decrease in the expression of Pfkm and Pfkl, but increased AMP/ADP ratio in BMDMs cultured with Se, along with other nucleotides that persisted even after LPS stimulation (Figure 3a). Together, such changes indicated functional glycolysis, albeit at low levels, necessitating auxiliary pathways such as PPP that utilize glycolytic intermediates to produce precursors of NADPH, along with increased selenoprotein expression, to maintain redox homeostasis as well as help buffer ROS to ultimately influence macrophage polarization (28).

Previous studies suggested that loss of Shpk (Carkl) led to a significant drop of NADH levels resulting in a redox shift (35). Compared to the Se-deficient cells, Se-supplementation of BMDMs held the NAD^+^/NADH ratio up to 4 h post LPS treatment, while NADP^+^/NADPH ratios were unchanged. GSH/GSSG ratio decreased temporally in Se supplemented macrophages upon LPS stimulation, but tightly maintained in Se-supplemented cells suggesting restoration of redox status in favor of resolution. Increased levels of Glu in Se-supplemented cells corroborated with restoration of GSH/GSSG balance and TCA cycle through increased α-ketoglutarate. Surprisingly, Txrnd1 and Hmox1 was mostly seen to increase transiently with Se suggesting a fine control by Nrf-2 dependent mechanisms.

Se treatment of BMDMs increased cellular levels of amino acids before and after LPS (up to 2 h), perhaps to support many functions of macrophages in addition to metabolic reprogramming. Though not exactly similar, a similar trend was reported earlier in whole liver (41). In addition, Se also increased monocarboxylate transporters, Slc16a1 and Slc16a3, and a high affinity copper transporter Slc31a1, to support increased pyruvate metabolism, TCA cycle and OXPHOS (data not shown). Macronutrients such as *myo*inositol (MI), a precursor for synthesis of phosphoinositides critical for signal transduction via second messengers IP3 and Ca^2+^, was consistently higher in the presence of Se, both in the absence and presence of LPS. Thus, a broader understanding of Se on MI metabolism in regulating macrophage functions may benefit from further studies.

Given the importance of fatty acid oxidation and OXPHOS, which drives the (M2) macrophage-mediated resolution and tissue repair (42), higher expression of Acsl3 and Acsl4, which are involved in fatty acid metabolism, along with Acadl, Acsl5, Adam17, Cyb5r3, Gcdh, Sgpp1, Sptlc1, Sptlc2, and the acetylCoA transporter slc33a1 (proteins represented by gene names), were temporally upregulated by Se (data not shown). In agreement with Se-dependent endogenous activation of PPARγ as reported previously (19), which is implicated in regulation of M2 genes, including Arg-1 and Mrc-1, fatty acid oxidation, adipocyte differentiation and glucose homeostasis, was upregulated upon LPS treatment (19, 43-45). Pathway network analysis suggested PPAR signaling associated proteins such as Csf2rb, Cbl, Akt3, Pik3r5, Stat3 and Irf9 were upregulated by Se supplementation. Stat3 mediates the anti-inflammatory effects of Il-10 (46) further reinforcing the anti-inflammatory role of Se via multiple mechanisms that promote the pro-resolution phenotype of BMDMs.

In conclusion, our studies demonstrate a key role for micronutrient Se in the form of selenoproteins in resolution, as seen in in-vitro and in-vivo models of inflammation. While the exact mechanistic underpinnings are still not clear, it is evident that exogenous addition of Se to Se-deficient BMDMs regulate key metabolic nodes, Sdh complex, Pkm, and Shpk (Carkl), in inflamed macrophages to promote anti-inflammation and resolution of inflammation (Figure 9). While these studies provide some explanation as to why Se-deficient sepsis patients respond better to Se therapy, our results further reiterate the importance of maintaining cellular Se, which tends to decrease in the elderly or those with chronic inflammatory disorders, to be important for effective resolution of inflammation.

**Figure 9.**
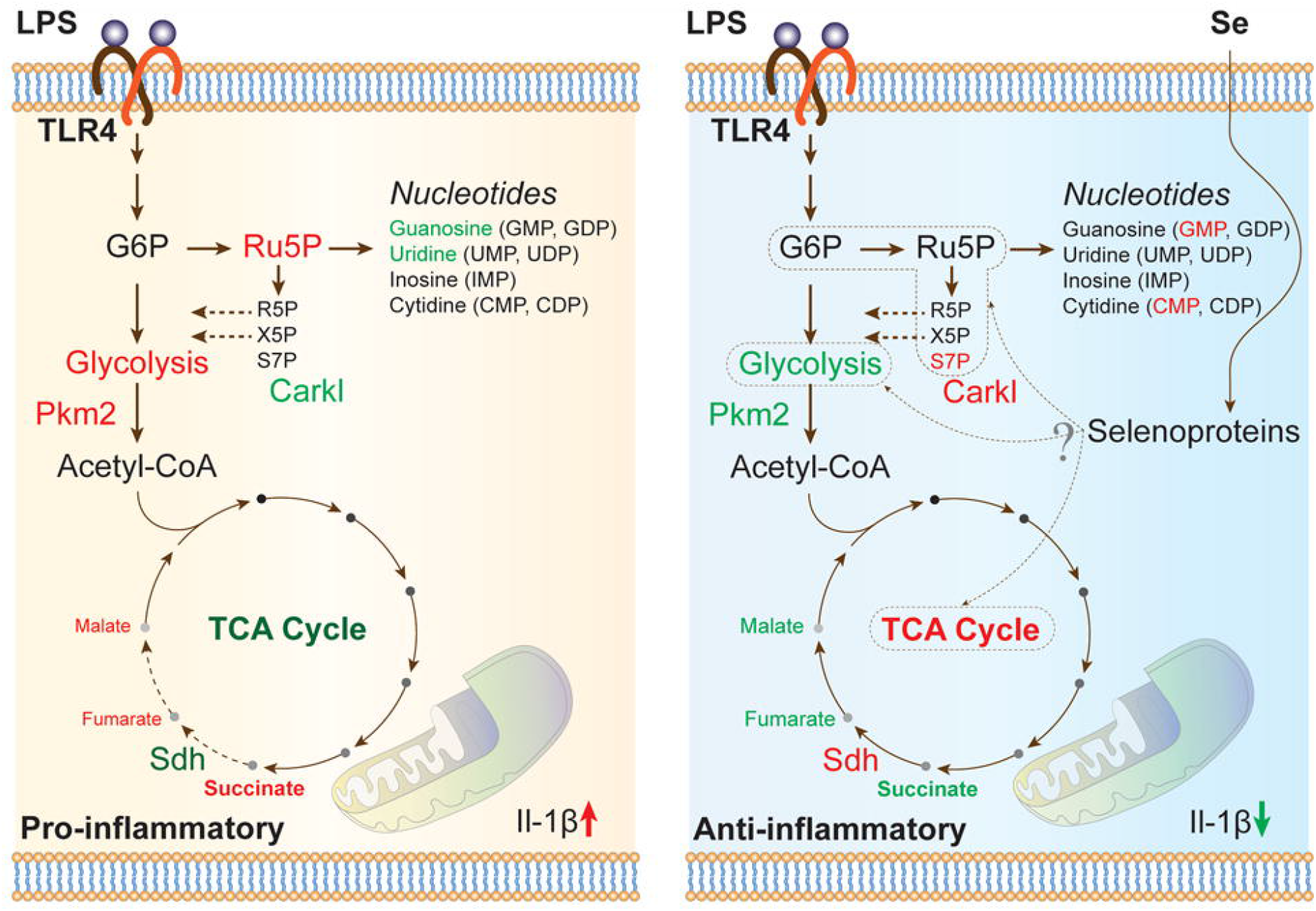
Schematic representation illustrating the effect of Se on BMDMs prior to and after LPS stimulation. Key metabolites and enzymes that were significantly upregulated (red) or downregulated (green) in LPS-treated (100 ng/ml) BMDMs ± Se (250 nM) are shown.

## METHODS

### Materials

TMT labeling reagent kit and Dulbecco’s Modified Eagle Media (DMEM) were procured from Thermo Fisher Scientific (NY, USA). Sodium selenite and lipopolysaccharide from *E. coli* (Serotype 0111:B4) were procured from Sigma Aldrich (St. Louis, MO, USA). Fetal bovine serum (FBS) and L929 fibroblasts were purchased from Atlanta biologicals (Atlanta, GA, USA), and American Type Culture Collection (Manassas, VA, USA), respectively. Basal levels of Se in the culture media was 7 nM as determined by ICP-MS-based methods. All other chemicals and reagents were of mass spectrometry grade.

### Mice and Genotyping

Three-week-old C57Bl/6 male mice were purchased from Taconic Biosciences, Inc. (Hudson, NY, USA) and maintained on either an AIN-76-based semi-purified Se-deficient diet (<0.01 ppm), or Se-supplemented diet (0.4 ppm) from Harlan-Teklad (Madison, WI, USA) for at least four weeks before being used in the experiments as previously described (21). A transgenic C57Bl/6 line carrying a lysozyme M Cre (*LysM^Cre^*) transgene was crossed to floxed *Trsp* (*Trsp^fl/fl^*) allele, both generously provided by Dr. Dolph Hatfield (NIH, MD, USA), as previously described (47). Genotyping was performed as previously described (21). All studies were preapproved by the Institutional Animal Care and Use Committee and the Institutional Biosafety Committee at The Pennsylvania State University, University Park, PA

### Bone Marrow-derived Macrophage Culture: LPS Stimulation

Three-week-old male C57Bl/6 mice were purchased from Taconic Biosciences, Inc. (Hudson, NY, USA) and maintained on either an AIN-76-based semi-purified Se-deficient diet (<0.01 ppm; Se-D) or Se-supplemented diet (0.4 ppm as selenite; Se-S) purchased from Harlan Teklad, (Madison, WI, USA) for 7 weeks before being used in the experiments. Femoral bone marrow from Se-D and Se-S mice were used as a source of BMDMs upon culture with no exogenous or 250 nM Se (as sodium selenite) addition, respectively, as described (26). Cells were cultured in biological triplicates for seven days with or without Se (as sodium selenite; 250 nM) as previously described (19, 26). On the eighth day, cells were stimulated with LPS (100 ng/ml; Sigma), where cells were treated with LPS for an hour followed by replacement with fresh culture media until harvest at 0, 4, 8, and 20 h post-stimulation. Cells were washed with ice cold sterile PBS, scraped, and cell pellets were stored at −80 °C until further processing.

### Murine Peritonitis and Flow Cytometry

Peritoneal inflammation was induced as described previously (48, 49). Se-S mice were treated intraperitoneally with dimethyl malonate (DMM) in PBS (160 mg/kg of body weight) or PBS alone three hours prior to administration of Zymosan A (10 mg/kg body weight) intraperitoneally. Mice were euthanized at 0, 12, 24, and 48 h post-Zymosan A injection, and peritoneal exudates were collected, centrifuged at 400 x *g* for 5 minutes at 4 °C, washed and resuspended in flow buffer (PBS containing 2% FBS; 1 mL), and 100,000 viable cells were analyzed by flow cytometry. Pellets were suspended in 100 μl of flow buffer and blocked with 0.25 μg Fc block (BD Pharmingen, CA, USA) for 10 min followed by staining with antibodies: 1 μl of PE-conjugated anti-mouse Ly-6G (BD Pharmingen, CA, USA), 1 μl of FITC-conjugated anti-mouse CD206 (Biolegend, CA, USA), and 10 μl of APC-conjugated anti-mouse F4/80 (Miltenyi Biotec, CA, USA) for 30 min at 4 °C in the dark. Cells were washed twice with flow buffer and centrifuged at 400 x *g* for 5 minutes at 4 °C and analyzed on a BD Accuri C6 Flow Cytometer and analyzed with FlowJo version 10 software (FlowJo, LLC, OR, USA). The resolution of inflammation defined as resolution index (R_i_) is the time taken to reduce the number of neutrophils at the site of inflammation by 50% (50). All animal protocols

### Sample Preparation for Proteomics, Data Acquisition and Analysis

The proteome was extracted from frozen cell pellets in RIPA buffer (Thermo Fisher Scientific, NY, USA) and protein concentration was determined by bicinchoninic acid (BCA) assay (Thermo Fisher Scientific, NY, USA). Equal amounts of protein (100 μg) was diluted with 100 mM triethyl ammonium bicarbonate buffer and reduced with dithiothreitol (10 mM) at 60 °C for 30 min, followed by alkylation with 2-chloroacetamide (65 mM) at room temperature in the dark for 30 min. The proteome was digested with proteomic grade trypsin at 1:50 (enzyme to substrate) ratio overnight at 37 °C. The peptide digest was labelled with TMT and fractionated (Lot number SH253273 and sample to TMT channel information is provided in Table S1). The proteomic dataset was acquired using a RSLC Ultimate 3000 nano-UPLC online coupled to an Orbitrap Fusion MS (Thermo Fisher Scientific, NY, USA). The proteomic dataset was acquired as previously described (26). Briefly, peptides were resolved on an Acclaim PepMap C18 column (2 microns, 75□μm i.d.□×□50□cm) with flow rate 300□nL/min using a 0.1% formic acid/acetonitrile gradient. For multinotch-MS3, the top ten precursors from each MS2 scan were fragmented by HCD followed by Orbitrap analysis (NCE 55; 60,000 resolution; AGC 5×104; max IT 120□ms, 100-500□m/z scan range). Mass spectral datasets were analyzed as previously described (26) using Proteome Discoverer (v2.2, Thermo Fisher Scientific, NY, USA) and searched against the SwissProt *Mus Musculus* database (17,424 proteins) using the following parameters: peptide and fragment mass tolerance were 10 ppm and 0.5 Da, respectively, with two miscleavages. The oxidation of methionine (15.995 Da) and deamidation of asparagine and glutamine (0.984 Da) were considered as variable modifications. TMT labeling of the N-termini of peptides, as well as lysine (229.163 Da) and cysteine carbamidomethylation (57.021 Da) were considered as static modifications. Relative quantitation using TMT reporter ions was performed using high-quality MS3 spectra. A percolator algorithm was used to determine the false discovery rate (FDR) and only peptides with an FDR□≤□0.01 were considered for further analysis.

Fold-change data were lowess normalized using StatsModels (statsmodels.org) and the correlations were measured using the Ordinary Least Squares linear regression method from SciPy (scipy.org). The sorted heatmap, stacked bar charts of the log2 fold-changes and related visualization was generated by using Matplotlib (matplotlib.org). Proteins were clustered using the k-means algorithm and separation analysis was performed using t-distributed Stochastic Neighbor Embedding algorithm from the scikit-learn machine learning library (scikit-learn.org). Cluster maps were generated using seaborn (seaborn.pydata.org). Clustering analysis compared protein’s abundance relative to naïve (untreated) 0 h, 0 Se sample and 4, 8, and 20 h following stimulation with LPS. In each of the experimental triplicates, two data points per protein were used for clustering. Values from the two replicates were averaged for drawing the curves in Figures 2a. Proteins identified in at least two replicates and with at least one unique peptide were included in this analysis. Kinetic deltagram algorithm was developed by Salis Lab at Penn State University (salislab.net) to visualize fold-change trends across different time points, as well as identify the window of most notable change, and was implemented in Python. Functional data analysis was carried using IPA (51), DAVID (52) and GSEA (53) for pathway enrichment.

### Sample Preparation for Metabolite Acquisition and Data Analysis

Metabolites were extracted from harvested BMDMs in prechilled 80% methanol. Samples were snap frozen in liquid nitrogen, vortexed and centrifuged for 20 minutes at 20000× *g* at 4 °C. The supernatant was dried using a SpeedVac and was resuspended in 3% aqueous methanol containing 100 μM chlorpropamide as internal standard. All samples were acquired in biological triplicate with randomized sample order. 10 μl aliquots of each sample were analyzed using LC-MS.

Metabolites were analyzed using reverse-phase UHPLC (C_18_ Hydro-RP column; Phenomenex) coupled to an Exactive Plus orbitrap MS (Thermo Fisher Scientific). A linear gradient from 3-100 (v/v) % methanol for 25 min was achieved with 3 % methanol, 10 mM tributylamine, 15 mM acetic acid (Solvent A) and 100% methanol (Solvent B) (54). Mass spectra were acquired in negative ion mode, with a scan range of 85-1,000 *m/z* with a resolution of 140,000 at *m/z* 200. A total of 47 metabolites were profiled across five time points of experiment (0, 2, 4, 8 and 20 h) in biological triplicate. Individual metabolite extracted-ion chromatograms were used to determine the integrated peak areas. All metabolite peak areas were normalized to the chlorpropamide and metabolite abundance was calculated relative to that derived from naive zero-hour samples. An in-house library generated from 292 authentic metabolite standards with experimentally observed accurate *m/z* values and retention times aided in the identification of metabolites (55).

### Western Immunoblot

Harvested cell pellets were resuspended in mammalian protein extraction reagent (Thermo Fisher Scientific, NY, USA) containing protease inhibitor mixture (Roche, Applied Science) and 5 mM sodium orthovanadate (Sigma, MO, USA) and incubated on ice for 20 min and vortexed for 10 min followed by centrifugation for 25 min at 20,000× *g* at 4 °C. Protein concentration was determined using BCA protein assay kit (Thermo Fisher Scientific, NY, USA). Proteins were resolved using a 12.5 % (% T) SDS-PAGE and transferred onto nitrocellulose membrane. The membrane was blocked with 7% (w/v) skim milk and probed with antibodies: anti-Sdha (Cell signaling; 1:2000), anti-SDHB (Proteintech; 1:4000), anti-Pkm (Cell Signaling; 1:10,000), anti-pPkm (Cell Signaling: 1: 10,000), anti-Shpk (Carkl) (MyBiosource; 1:2500) and anti-β-actin (Fitzgerald; 1: 25000). Data from three independent experiments were used in densitometric analysis, while only a representative Western blot is shown for brevity.

## Data availability

The mass spectrometry proteomics data generated in this study have been deposited at the ProteomeXchange Consortium via the PRIDE (56) partner repository with the dataset identifier PXD023005.

## Acknowledgements

We thank current and past members of the Prabhu laboratory for their invaluable suggestions and Dr. Sougat Misra for his timely help with the artwork.

## Author contributions

AMK, AES, and KSP conceived the project, designed research, analyzed data, and wrote the manuscript; AMK performed all the experiments, AH and HMS developed algorithms and performed statistical analysis and visualization of proteomic and metabolomic datasets, TJL performed western blot experiments, VB and KC digested the proteome, labeled, and acquired the dataset, PBS and ADP supervised acquisition of metabolomics data, BA created *Trsp* KO mice and assisted with manuscript preparation.

## Funding and additional information

These studies were funded, in part, by grants from the National Institutes of Health and Office of Dietary Supplements NIH DK077152, USDA-NIFA Hatch Project USDA-NIFA (Project # 4605; Accession # 1010021), Penn State Cancer Institute, The Metabolomics Facility and the Defense Advanced Research Projects Agency (FA8750-17-C-0254). The content is solely the responsibility of the authors and does not necessarily represent the official views of the National Institutes of Health.

## Abbreviations

Se: Selenium
Trsp: tRNA^[Ser]Sec^
TMT: tandem mass tag
SDH: succinate dehydrogenase complex
PKM: pyruvate kinase
SHK: sedoheptulosekinase
LPS: lipopolysaccharide
IL-4: interleukin-4
PPP: pentose phosphate pathway
OXPHOS: oxidative phosphorylation
TCA: tricarboxylic acid
DMM: dimethylmalonate
BMDMs: bone marrow-derived macrophages

## SUPPORTING INFORMATION

**Figure S1. Effect of LPS on the expression of prototypical genes in BMDMs.** LPS-dependent expression of prototypical proteins in BMDMs by proteomic studies post 20 h.

Fold changes in expression were calculated relative to unstimulated cells. For these studies, BMDMs cultured in the absence of Se were used. Mean of n= 3 independent experiments per time point were used.

**Figure S2. Clustering and functional analysis of Se-dependent effects of LPS in BMDMs.** Protein cluster enrichment 1 to 7 enrichment of post-LPS proteome and their regulation kinetics. In column **a** represent post-LPS proteome and their expression kinetics is depicted in columns **b** and **c**. The red and blue colors in columns **b** and **c** depict modulation with Se and without Se, respectively. The black patches in column **c** highlights the time interval showing most notable changes in kinetics upon stimulation with LPS. Functional analysis depicting specific metabolic pathways enriched within each cluster is shown in column **d**.

**Figure S3. Modulation of the mitochondrial proteome changes by Se supplementation (as sodium selenite, 250 nM) positively impacts the TCA cycle and OXPHOS related proteins. a.** The mitochondrial proteome was classified into six clusters based on the kinetic regulation of protein expression upon LPS stimulation. Kinetic regulation are depicted in column **b** and **c**. Pathway analysis identified the specific metabolic pathways enriched within respective cluster are depicted in column **d**.

**Figure S4. Differential modulation of cytosolic ROS (a) and mitochondrial ROS (b) in BMDMs by Se supplementation following LPS stimulation.** BMDMs were isolated from WT mice and cultured in media with or without the presence of selenite (250 nM). BMDMs were stimulated with 100ng/ml LPS. After 1hr stimulation, LPS containing media were removed and replaced with fresh media and cells were continuously cultured for indicated time periods. Following stimulation, BMDMs were then stained for 30 min with CellROX Deep Red (500 nM) flow cytometry stain for cytosolic ROS (**panel A**) or MitoSOX Red stain (5 uM) for mitochondrial ROS (**panel B**), followed by 15 mins of the dead cell staining with propidium iodide (PI) or 7-aminoactinomycin D (7-AAD), respectively. Data shown are n= 3 per group ± SEM; *p<0.05.

**Figure S5. Selenoprotein-dependent changes in glycolytic metabolites in WT and *Trsp^fl/fl^ LysM^Cre^* BMDMs.** BMDMs isolated from WT and *Trsp^fl/fl^ LysM^Cre^* mice were stimulated for 2-20 h with LPS. Metabolites were isolated and subjected to targeted metabolomics. Temporal regulation of glycolytic metabolites was analyzed in BMDMs. Phosphoenol pyruvate and pyruvate, was observed to be distinctly different in Se supplemented WT when compared to *Trsp*^fl/fl^ *LysM*^Cre^ BMDMs suggesting a causal association with selenoproteins expression. For each time point upon LPS stimulation, mean of three independent experiments (± SEM) were considered for each genotype. * p<0.0001, ** p<0.001, * p<0.01

**Figure S6. Selenoprotein-dependent changes in PPP metabolites in WT and *Trsp^fl/fl^ LysM*^Cre^ BMDMs post LPS.** Temporal regulation of PPP metabolites, especially sedoheptulose-1/7-phosphate was observed to be distinctly different in Se supplemented WT BMDMs compared to *Trsp*^fl/fl^ *LysM*^Cre^ BMDMs suggesting their modulation to be causally associated with selenoprotein expression. For each time point upon LPS stimulation, mean of three independent experiments (± SEM) were considered for each genotype. * p<0.0001, ** p<0.001, * p<0.01

