## Supplementary Data for "Selenium-dependent metabolic reprogramming during inflammation and resolution"

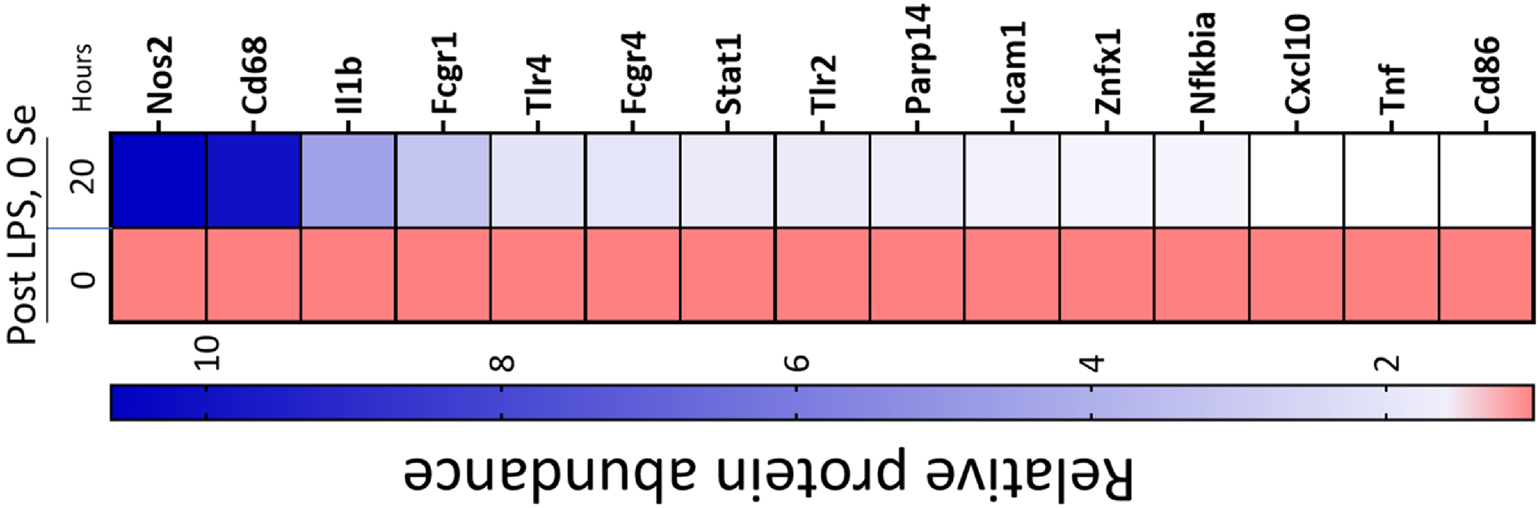


Supplementary Figure S1

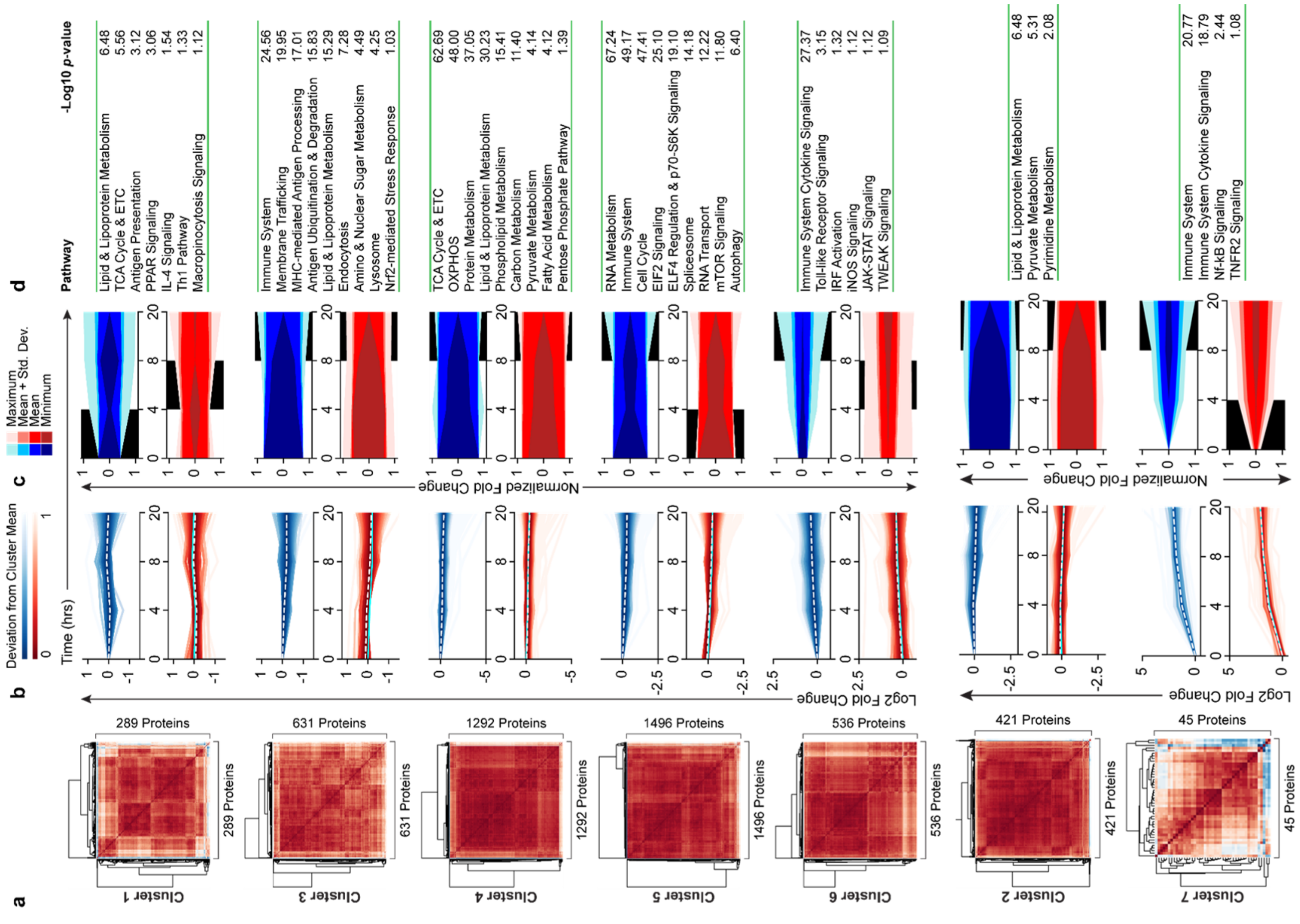


Supplementary Figure S2

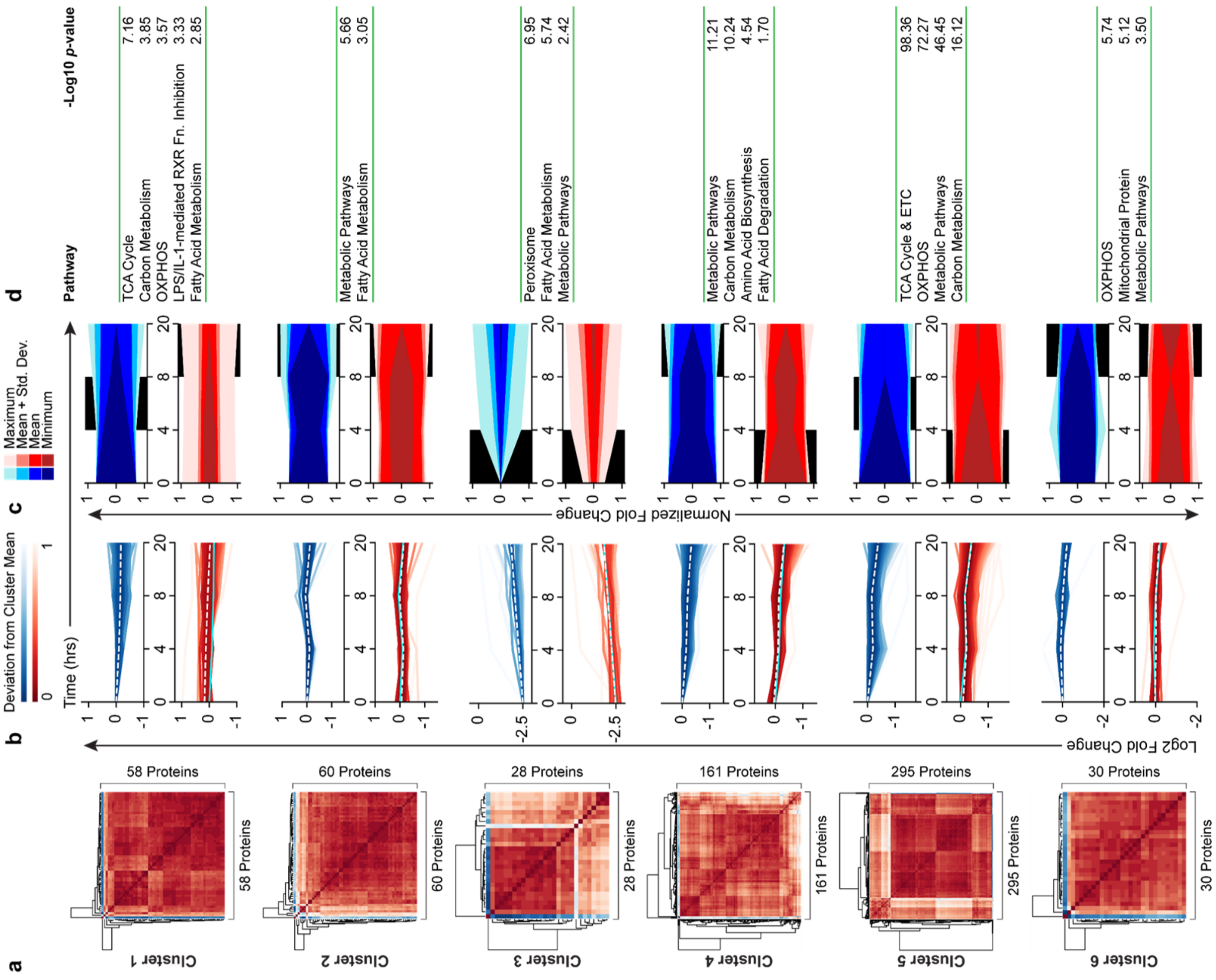


Supplementary Figure S3

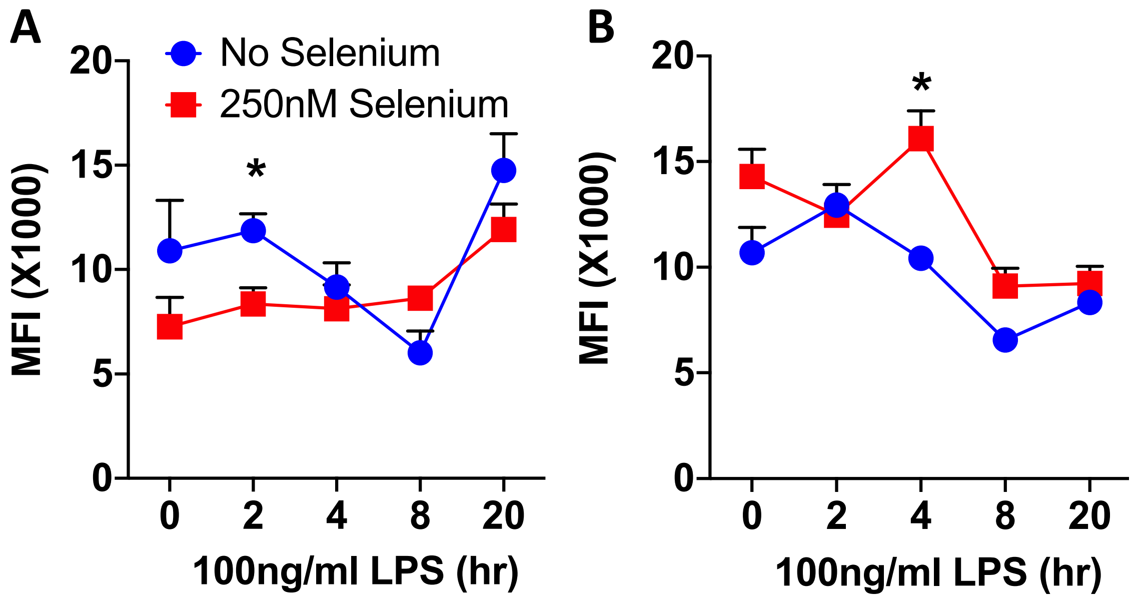


Supplementary Figure S4

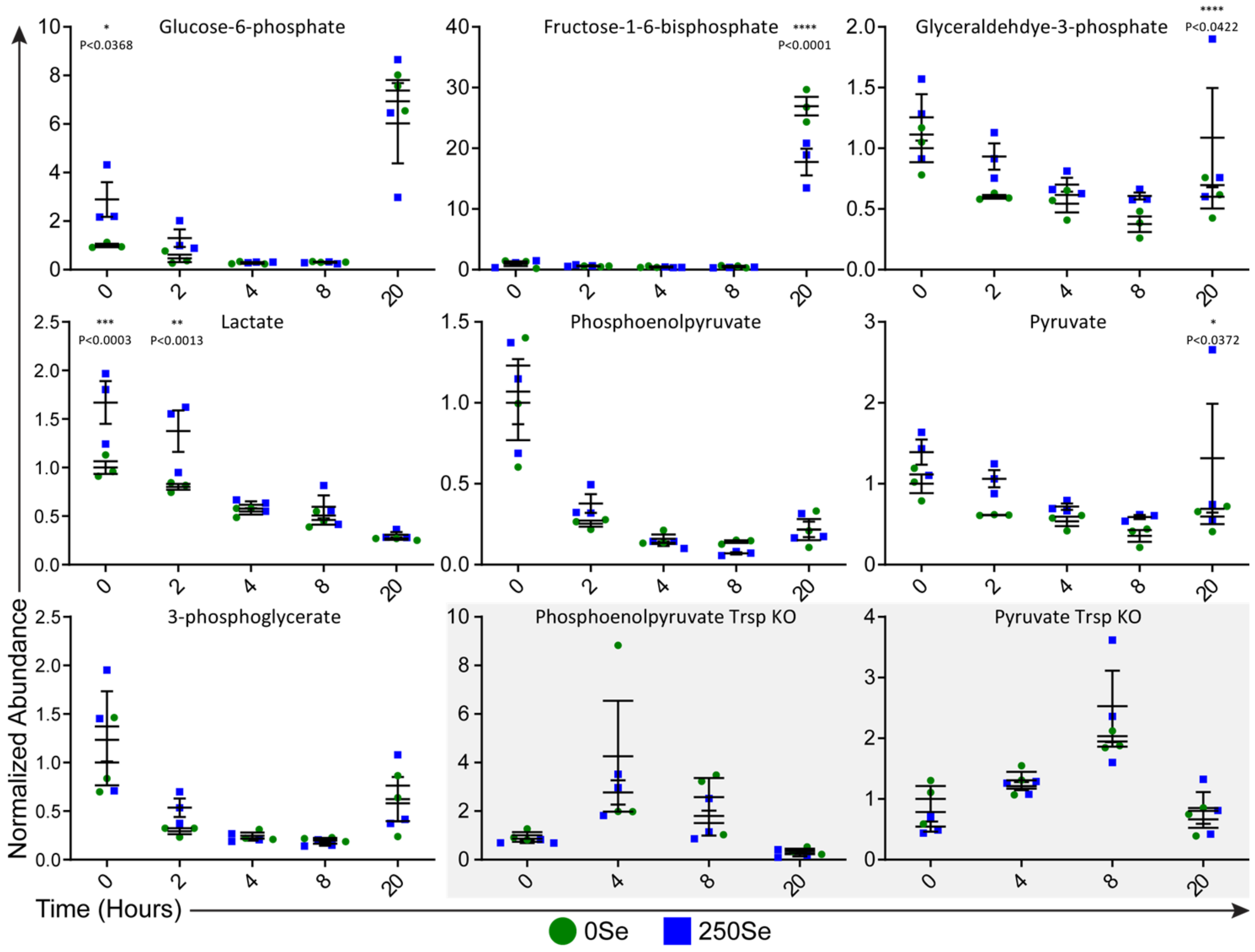


Supplementary Figure S5

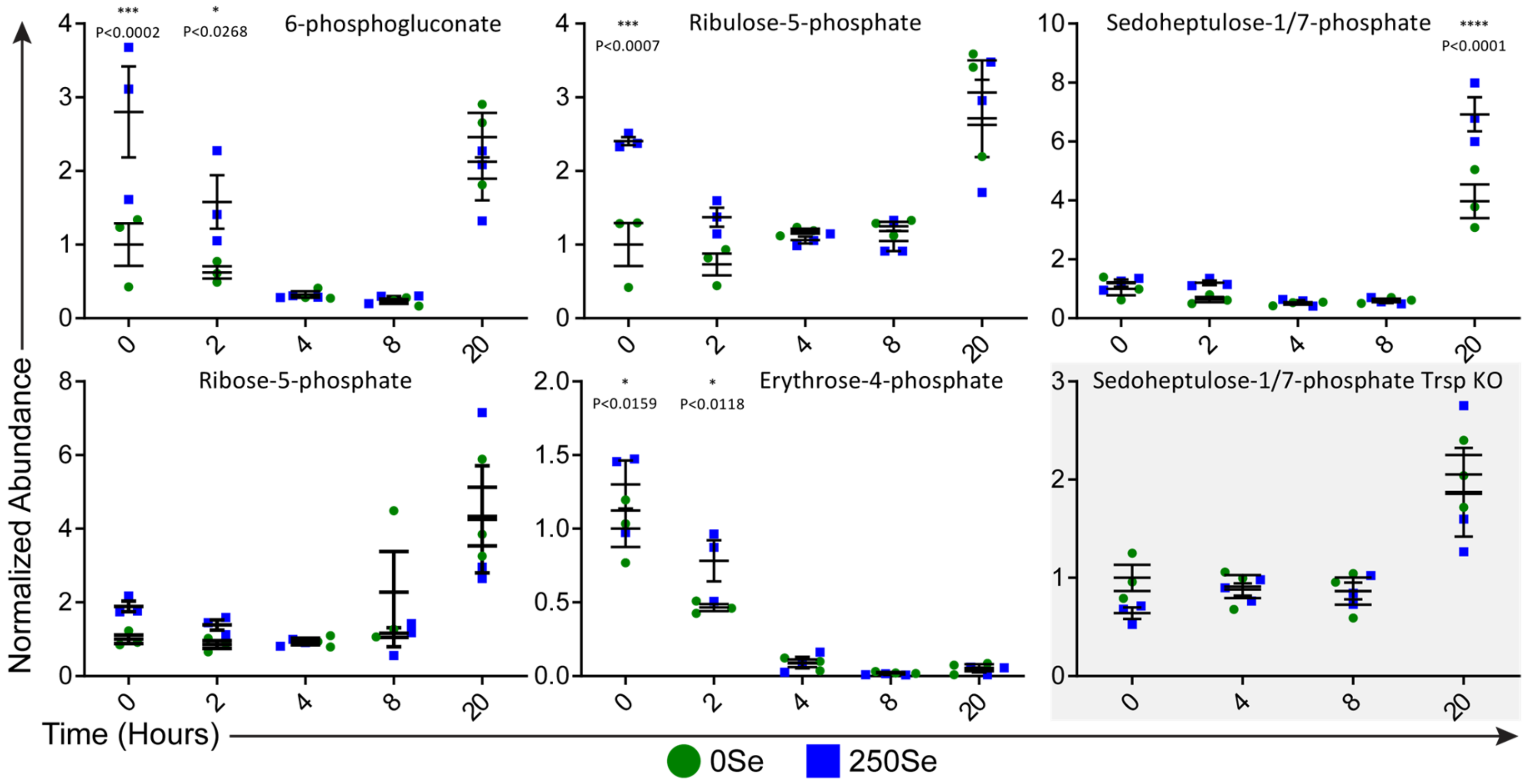


Supplementary Figure S6
